# Asian Zika virus isolate significantly changes the transcriptional profile and alternative RNA splicing events in a neuroblastoma cell line

**DOI:** 10.1101/660209

**Authors:** Gaston Bonenfant, Ryan Meng, Carl Shotwell, J. Andrew Berglund, Cara T. Pager

## Abstract

Alternative splicing of pre-mRNAs expands a single genetic blueprint to encode multiple functionally diverse protein isoforms. Viruses have previously been shown to interact with, depend on, and alter host splicing machinery. The consequences however incited by viral infection on the global alternative slicing (AS) landscape are under appreciated. Here we investigated the transcriptional and alternative splicing profile of neuronal cells infected with a contemporary Puerto Rican Zika virus (ZIKV^PR^) isolate, the prototypical Ugandan ZIKV (ZIKV^MR^) isolate and dengue virus 2 (DENV2). Our analyses revealed that ZIKV^PR^ induced significantly more differential changes in expressed genes compared to ZIKV^MR^ or DENV2, despite all three viruses showing equivalent infectivity and viral RNA levels. Consistent with the transcriptional profile, ZIKV^PR^ induced a higher number of alternative splicing events compared to ZIKV^MR^ or DENV2, and gene ontology analyses highlighted alternative splicing changes in genes associated with mRNA splicing. All three viruses modulated alternative splicing with ZIKV^PR^ having the largest impact on splicing. ZIKV alteration of the transcriptomic landscape during infection caused changes in cellular RNA homeostasis, which might dysregulate neurodevelopment and function leading to neuropathologies such as microcephaly and Guillain-Barré syndrome associated with the ZIKV infection.

## Introduction

Zika virus (ZIKV) is a re-emerging mosquito-borne flavivirus that is classified within the *Flaviviridae* family. Other notable flaviviruses include Dengue virus (DENV), Yellow Fever virus (YFV), West Nile virus (WNV) and Tick-borne encephalitis virus, all of which are primarily transmitted via the bite of an infected mosquito or tick [1]. Flavivirus infections rarely result in death and common symptoms include maculopapular rash, fever, and achy joints [2]. Until the early 2000s, only 13 confirmed ZIKV infections in humans were reported [3–6]. The first major outbreak of ZIKV occurred in 2007 on Yap Island [7], followed by a 2010 outbreak in Cambodia [8], and an outbreak in French Polynesia in 2013 which resulted in more than 29,000 human infections [9]. This Asian lineage of ZIKV expanded west and in 2015 efforts were redirected towards understanding the link between ZIKV infection and the associated neurological pathologies that are now termed Congenital Zika Syndrome (CZS) [10,11]. To date there are no antivirals or a licensed vaccine to prevent ZIKV infection. Therefore, to develop effective therapies and thus limit the dreadful symptoms associated with ZIKV infection it is critical to understand viral-host interactions and ZIKV pathogenesis.

The striking feature of the recent ZIKV outbreak in the America’s was the correlation between intrauterine ZIKV infection and the devastating consequences to fetal brain development resulting in microcephaly, cortical malformations and intracranial calcifications [12–15] and the increased number of cases of Guillain-Barré syndrome in adults [16–19]. As a first step to elucidating ZIKV-directed mechanisms resulting in neurological anomalies *in vitro, ex vivo*, primary cells and *in vivo* mouse infection studies were undertaken. These studies determined that ZIKV infects neuroepithelial stem cells and radial glia cells, resulting in cell cycle arrest, altered differentiation, increased cell death, and affected the thickness of neuronal layers [20–24]. These outcomes at the cellular level were the result of ZIKV disrupting centrosomes, changing the cell division plane, inducing apoptosis and altering signaling pathways [12,20–23,25,26]. At the genetic level, ZIKV was shown to dysregulate the transcription of cell-cycle, DNA repair, immune response, cell death and microcephaly genes [12,20–23,25,26]. Infection of neural stem cells and other neuronal cell lines by the original ZIKV strain isolated in Uganda in 1947 versus the Asian isolates, including those isolated from the 2015 outbreak in the Americas report differences in infectivity. Despite these reported infectivity differences, RNA-seq studies showed that the changes in the transcriptome were less dramatic [12], suggesting that transcriptional changes alone do not entirely explain the ZIKV neuropathology.

We recently showed that during ZIKV infection HuR (or ELAVL1) is re-localized from the nucleus to ZIKV replication sites [27]. Since the ELAVL family of proteins regulate mRNA splicing and stability [28,29], we posited that the localization of certain RNA-binding proteins, such as HuR, could impact RNA transcription as well as mRNA splicing and stability and thus contribute to the misregulation of cellular pathways critical for neuronal development. Indeed, molecular diversity within the central nervous system is in part the result of alternative splicing events [30]. Studies of developing cortices in primates [31] and rodents [32] showed variation of alternative exons and brain- or neuron-specific splicing patterns changed dramatically [33,34]. Moreover, the temporal and cell-type specific regulation of AS events was in large part due to recognition of regulatory sequences within pre-mRNA transcripts by RNA-binding proteins (RBPs) enriched in neurons, such as Rbfox and neuronal ELAV proteins [30].

In this study, we used RNA-seq to investigate transcriptional profiles and alternative splicing events in a neuroblastoma cell line following infection with a modern isolate of ZIKV circulating in the Americas (PRVABC59; ZIKV^PR^), the original 1947 ZIKV isolate from Uganda (MR766; ZIKV^MR^) and dengue virus serotype 2 (DENV2). Analysis of global transcription revealed seven times more changes in gene expression following infection with ZIKV^PR^ compared to infection with ZIKV^MR^ or DENV2. Moreover, the number of alternative splicing events correlated with the transcriptional profile of each virus infection, where the skipping of exons exceeded other alternative splicing events. Our study establishes the foundation to further investigate the impact of specific alternatively spliced genes on neuronal development and subsequent neuropathologies observed following ZIKV infection.

## Materials and Methods

### Cell Maintenance

SH-SY5Y neuroblastoma cells (ATCC CRL-2266) were cultured at 37°C with 5% CO_2_ and maintained in Eagle’s minimum essential medium (Sigma), F-12 Ham with NaHCO_3_ (Sigma), and 10% fetal bovine serum (FBS; Seradigm). *Aedes albopictus* cells (C6/36; ATCC CRL-1660) were grown at 27°C supplemented with 5% CO_2_ and maintained in Eagle’s minimal essential medium (Sigma) supplemented with 10% FBS, sodium pyruvate (0.055 g/L; Life Technologies), Fungizone (125 μg/L; Life Technologies), and penicillin and streptomycin (50,000 units/L penicillin, 0.05 g/L streptomycin; Life Technologies).

### Preparation of Virus Stocks

ZIKV^MR^ (Uganda MR766 strain) was a gift from Dr. Brett Lindenbach (Yale University) and ZIKV^PR^ (Puerto Rico PRVABC59 strain) was a gift from Dr. Laura Kramer (Wadsworth Center NYDOH) and the CDC. To create virus stocks, C6/36 cells were seeded into 15 cm tissue culture plates, and cells were infected near confluence. In particular, a viral master mix was prepared (5 mL PBS and 500 μL virus aliquot per plate), media was aspirated from each plate, and the viral master mix was added to cells. The plate was returned to the incubator for one hour with rocking every 15 minutes. After the one-hour incubation, 10 mL complete media was added, and the plate was returned to incubate for seven days. After seven days, media was transferred to 50 mL conical tubes, centrifuged at 1,000xg for 5 minutes, and 500 μL aliquot of the viruses were prepared and stored at −80°C. To ensure successful infection, 3 mL TRIzol reagent was added to the culture plate and RNA was extracted for RT-qPCR and RNA-seq analysis. Virus titers were determined by plaque assay as previously described [27]. Aliquots of Dengue virus (DENV) serotype 2 (OBS-2629) isolated in Peru in 1996 were gifted to the Pager laboratory by Dr. Alexander Ciota (Wadsworth Center NYDOH).

### Viral Infections

SH-SY5Y cells were seeded in 6 cm tissue culture plates with 3 mL complete media. When cells neared 90% confluence, a control plate of cells was counted to determine multiplicity of infection (moi). SH-SY5Y cells were infected at a moi of 5. In particular, the appropriate amount of virus was diluted in PBS to 1.5 mL total volume which was then added to each plate. Plates were returned to the incubator for one hour with rocking every 15 minutes, after which 1.5 mL complete media was added. Cells were harvested at one-day post-infection.

### Harvest of Virus-Infected Cells

Cells were harvested by first aspirating media from the tissue culture plates, gently washing the cells with 1 mL PBS, and lysing the cells in 500 μL TRIzol reagent (Life Technologies). The cells in TRIzol were manually scraped, transferred to a 1.5 mL tube, and RNA was extracted following manufacturers’ recommendations. Following ethanol cleanup, RNA was resuspended in 20 μL Ambion 0.2 μm filtered water.

### Immunofluorescence Analysis

SH-SY5Y cells were seeded into 24-well plates treated with poly-D-lysine (1 mg/mL; Sigma). When cells reached 90% confluence, 2 wells were trypsinized and counted to determine moi. Similar to infection in the 6 cm tissue culture plates, SH-SY5Y cells were infected at a moi of 5 in a total viral master mix volume of 300 μL. One hour after the addition of virus, 600 μL complete media was added to each well. One day post-infection, cells were washed twice with PBS, fixed with 4% paraformaldehyde in PBS for 10 minutes at room temperature, and then permeabilized with 100% iced methanol for 10 minutes. Cells were washed in blocking buffer (PBS-1% fish gelatin (FG); Sigma) three times for 15 minutes. Mouse-anti-J2 dsRNA antibody (Scicons) was diluted in blocking buffer (1:250 v/v), added to the appropriate wells, and incubated overnight at 4°C. Donkey-anti-mouse-488 antibodies (Life Technologies) diluted in blocking buffer (1:200 v/v) were added for 1 hour at room temperature in the dark. Hoechst-33342 (Life Technologies) was applied for 15 minutes. Before and after application of antibodies, cells were washed with blocking buffer three times for 15 minutes. Finally, the cells were washed twice with PBS for 5 minutes, and the cells were visualized on the Evos FL cell imaging system (ThermoFisher Scientific).

### RT-qPCR

To confirm ZIKV and DENV infection prior to preparation of RNA-seq libraries, qRT-PCR was performed with virus-specific primers (Table S1), and the amount of viral determined relative to β-actin mRNA levels. RNA for each sample (100ng) and 10 μM forward and reverse primer were added to RT-PCR master mix containing RT enhancer, Hot-Start master mix, and Verso Enzyme mix (Life Technologies). Reactions were carried out beginning with reverse transcription at 50°C for 15 minutes. The RT enzyme was heat-inactivated at 95°C for 5 minutes followed by 35 cycles of 20 second denaturation at 95°C, annealing at 60°C for 30 seconds, and extension at 72°C for 1 minute.

### Preparation of Libraries for RNA-seq

For RNA-seq analysis libraries were prepared using the NEBNext® poly(A) magnetic isolation module (NEB E7490), NEBNext® Ultra™ Directional RNA library prep kit for Illumina® (NEB E7760) and NEBNext® Multiplex oligos for Illumina® (NEB E7335S). There libraries were concurrently prepared from three independent experiments of mock-, ZIKV (MR766 and PRVABC) and DENV-2-infected SH-SY5Y cells. The Poly(A) mRNA was isolated via magnetic beads (NEB #E7490) from RNA isolated by TRIzol-extraction. In brief, RNA (100 ng) was diluted in nuclease-free water to a final volume of 50 μL and oligo dT beads in 50 μL RNA binding buffer was added. Following a 65°C incubation for 5 minutes, samples were cooled to 4°C, left at room temperature for 5 minutes, and washed. Next, 50 μL Tris buffer was added to each sample, mixed, and incubated at 80°C for 2 minutes. After cooling to room temperature, 50 μL RNA binding buffer was added and incubated for 5 minutes. Beads were washed, supernatant was aspirated, and mRNA was eluted by the addition of 11.5 μL First Strand Synthesis Reaction Buffer and Random Primer Mix. Samples were placed in a thermal cycler set to 94°C for 15 minutes. Fragmented RNA was transferred to a new tube and placed on ice.

First strand cDNA synthesis reaction was assembled according to manufacturers’ specifications and added to samples, which were placed in a thermal cycler set to 25°C for 10 minutes, 42°C for 15 minutes, 70°C for 15 minutes, and a 4°C hold. Second strand synthesis was assembled as instructed by manufacturer and samples were incubated for one hour at 65°C. Double-stranded cDNA was purified using SPRIselect Beads. Briefly, cDNA incubated with beads at room temperature for 5 minutes, supernatant was aspirated, and the beads were washed three times with freshly prepared 80% ethanol. After the final wash, beads were air-dried and DNA was eluted by adding 53 μL 0.1X TE buffer, vortexed, incubated at room temperature for 2 minutes, and the supernatant was transferred to a new tube.

Adaptor ligation was performed using NEBNext Ultra II End Prep Reaction according to NEB’s instructions. NEBNext adaptors diluted 5-fold in iced adaptor dilution buffer, ligation enhancer, and ligation master mix were added to each tube, gently mixed, and incubated at 20°C for 15 minutes. Following the 15-minute incubation, USER Enzyme was added to each reaction, which was incubated at 37°C for 15 minutes. Ligation reactions were purified using SPRIselect beads as described above, though DNA was eluted in 17 μL 0.1X TE buffer. Eluted DNA was transferred to a new tube and PCR enrichment of the adaptor ligated DNA was performed using NEBNext Multiplex Oligos for Illumina (Set 1, NEB #E7335). The PCR reaction mix was assembled according to manufacturers’ recommendations and placed in a thermal cycler set to the recommended conditions. Based on the amount of input material, 10 cycles were carried out. PCR reactions were purified using SPRIselect beads, eluted in 23 μL 0.1X TE buffer, and transferred to a clean tube. Prior to sequencing, libraries were checked for purity via bioanalyzer and RT-PCR. RNA-seq was performed using the Illumina NextSeq500.

### Bioinformatics preprocessing

Raw sequencing reads were analyzed for quality using FASTQC [35]. Following quality and control analysis, raw sequencing reads were aligned back to the hg19 reference genome using STAR [36] for AS analysis. Post-alignment, BAM files were indexed using samtools [35]. Count tables were generated with Salmon [37] for differential gene expression (DGE) analysis.

### Differential Gene Expression Analysis and Alternative Splicing

DGE analysis was performed using DESeq2 [38]. Genes that exhibited an adjusted P value of less than or equal to 0.05 were deemed statistically significant. Gene enrichment analysis of the statistically significant differential gene expression was done with Panther [39]. Heatmaps and Volcano plots were produced using the Enhanced Volcano [40] and pheatmap [41] packages, respectively, in R [42].

Differences in splicing between the individual libraries were assessed using replicate multivariate analysis of transcript splicing (rMATS) [43]. Splicing events were filtered using a custom Python [44] script utilizing a cutoff of an absolute value of change in <u>P</u>ercent <u>S</u>pliced-<u>I</u>n (PSI) of greater than or equal to 0.10 and a false-discovery rate (FDR) of less than or equal to 0.05. Individual events were visualized using Sashimi plots [45] utilizing the rmats2sashimiplot tool (http://www.mimg.ucla.edu/faculty/xing/rmats2sashimiplot/).

### Alternative Splicing Analyses

Following cell infection and TRIzol extraction, RNA concentration was determined using a Nanodrop and 500ng of each sample was used in a reverse transcription with SuperScript IV using random hexamer primers (IDT). Half the suggested amount of Superscript was used in these reactions. The cDNA then underwent PCR for 30 cycles using NEB’s 2X Taq Master Mix and the primer sets listed in (Table 1). The resulting products were then run with a Fragment Analyzer capillary electrophoresis using the DNF905 1-500 base pair kit (Agilent). Quantification was performed using the following equation: Percent spliced in (PSI) = (RFU of Inclusion Band)/((RFU Inclusion Band+RFU Exclusion Band))×100%, where RFU represents Relative Fluorescence Units. Graphs were made using GraphPad Prism software and unpaired t tests were performed in the program to show significance.

### Statistics

For Figure S2A–E, reads were aligned to the hg38 genome using HISAT2, followed by quantification of features using FeatureCounts. Transcript counts were used for analysis via Limma-voom with no gene annotation file. Low expressed genes were filtered out based off counts per million (cpm) values < 1.0 or samples containing 0 genes. Genes were filtered based on a minimum log2 fold change of 0.5 with a *P*-value adjusted threshold [46] of 0.05. Finally, genes were normalized using the TMM method. Job-dependencies used and versions include Bioconductor-limma (Version 3.34.9), bioconducter-edge (Version 3.20.7), and r-statmod (Version 1.4.30).

### Data Access

Sequencing data from RNA-Seq were deposited to the NCBI GEO and are available under accession number GEO:PENDING

## Results

### SH-SY5Y cells are permissive for both Zika virus isolates and Dengue virus

Phylogenetic analyses reveal an African and Asian ZIKV lineage [47,48]. Neurological disorders, such as congenital Zika syndrome in newborns and Guillain-Barré syndrome in adults, have been causally linked to the Asian lineage ZIKV strain [49]. In this study our goal was to investigate the consequence of ZIKV infection on alternative splicing and link such changes to the neuropathologies associated with ZIKV infection. To this end we surveyed the transcriptional and alternative splicing (AS) landscape induced by ZIKV infection in SH-SY5Y, a neuroblastoma cell line. Because we were interested in differences in gene expression and alternative splicing resulting from infection with a modern ZIKV strain isolated in Puerto Rico in 2015 (PRVABC-59; ZIKV^PR^) [50], we also included the original ZIKV isolate from Uganda (MR766; ZIKV^MR^) [51] for comparison as well as a flavivirus that is not known cause neuropathies namely a serotype 2 dengue virus isolate from Peru (DENV2).

We first examined the permissiveness of SH-SY5Y cells to ZIKV^PR^, ZIKV^MR^ and DENV2-infection. In particular, SH-SY5Y cells were infected at a moi of 5, and at 24 hours post-infection we examined virus infection (Figure 1). Specifically, we determined the extent of infection by fixing and processing cells for immunofluorescence analysis. Using an antibody that specifically detected double-stranded RNA, an intermediate of flavivirus replication (Figure 1A), we observed between 31% and 40% of all cells were infected with ZIKV^PR^, ZIKV^MR^ and DENV2 (Figure 1B). We also isolated total RNA and determined the relative abundance of viral RNA by RT-qPCR. To this end we used primers specific to ZIKV^PR^, ZIKV^MR^ and DENV2 and normalized the Ct values to those obtained for β-actin mRNA in each sample. RT-qPCR revealed no statistically significant difference in the viral RNA abundance (Figure 1C). Together these data indicate that ZIKV^PR^, ZIKV^MR^ and DENV2 show similar infection levels and gene expression in SH-SY5Y cells. Moreover, these data indicate SH-SY5Y cells are equally permissive for flavivirus infection and thus presented an unbiased analysis of transcriptome changes and alternative splicing events resulting from viral infection.

**Figure 1:**
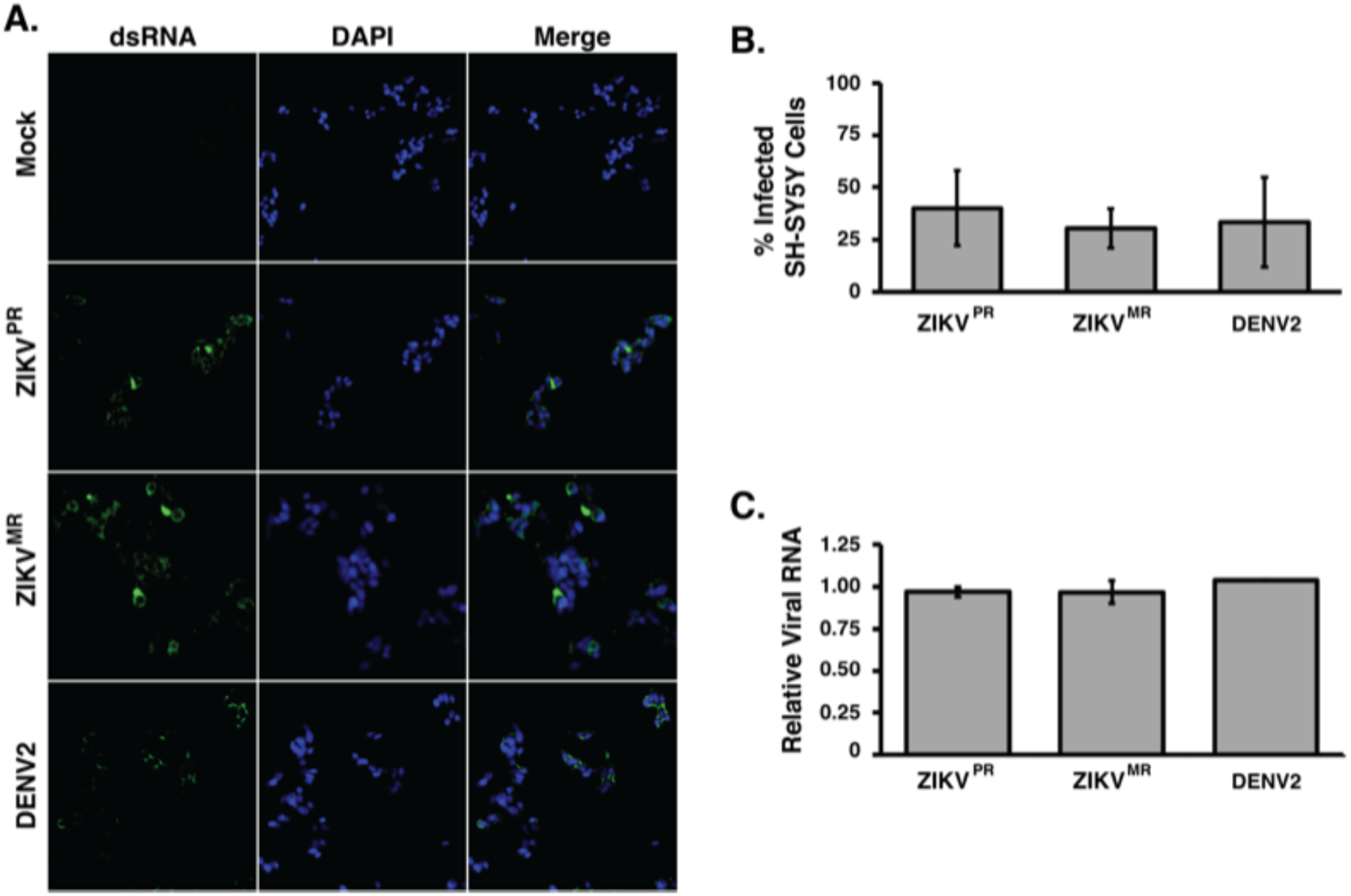
SH-SY5Y cells are permissive to ZIKV and DENV2 infection. A) Immunofluorescence images of SH-SY5Y cells infected at a moi of 5 and fixed 1 day post-infection. The cells were stained for dsRNA to detect virus infected cells, and Hoechst to visualize the nuclei in the field of view. B) Quantification of percent infected cells. SH-SY5Y cells in 48-well plates seeded were infected at a moi of 5, fixed 1 dpi, and stained for dsRNA and Hoechst. A 20x objective was used to capture three images per well were taken such that for each virus for a total of more than 400 cells for each replicate was counted. The percent infected cells were determined. At least three biological replicates were performed. C) RT-qPCR analysis of SH-SY5Y cells infected at a moi of 5. Primers targeting the coding regions of each virus were used along with primers for β-actin mRNA. Relative viral RNA levels were calculated by standardizing relative fluorescent units at Cq for each virus against β-actin mRNA. Error bars represent standard deviation. No significant difference was determined for the number of cells counted in panel 1B, or the relative abundance of viral RNA in panel 1C.

### ZIKV^PR^ significantly alters differential gene expression compared to ZIKV^MR^ and DENV2

To compare changes in the transcriptome we examined poly(A)-selected transcripts isolated from mock- or virus-infected SH-SY5Y cells 24 hours post-infection and performed RNA-seq analysis on the Illumina platform (Figure 2A). Prior to library preparation, we confirmed virus infection by RT-qPCR (Figure 1C). The analyses were derived from 75-nt paired end reads from three biological replicates. We obtained a total of 35-48 million reads from the three independent experiments which were mapped back to the reference human genome. Transcript expression was quantified using Salmon [37] and differential expression (DE) analysis was performed using DESeq2 [38] (Figure 2A). We observed a significant difference in DE expression resulting from ZIKV^PR^ infection indicating the modern isolate of ZIKV dramatically changed the transcriptome compared to infections with the African isolate ZIKV^MR^ and DENV2 (Figure 2B). A total of 1485 genes were differentially expressed in cells infected with ZIKV^PR^, nearly 6- and 8-times fold higher compared to ZIKV^MR^ and DENV2-induced DE genes, respectively (Figure 2B). Of these, 703 and 651 genes were up- and downregulated respectively (Figure S1). Thirty-three DE genes were common to the three flaviviruses, while 102 DE genes were shared between ZIKV^PR^ and DENV2, and 116 DE genes were common between the two ZIKV isolates (Figure 2B).

**Figure 2:**
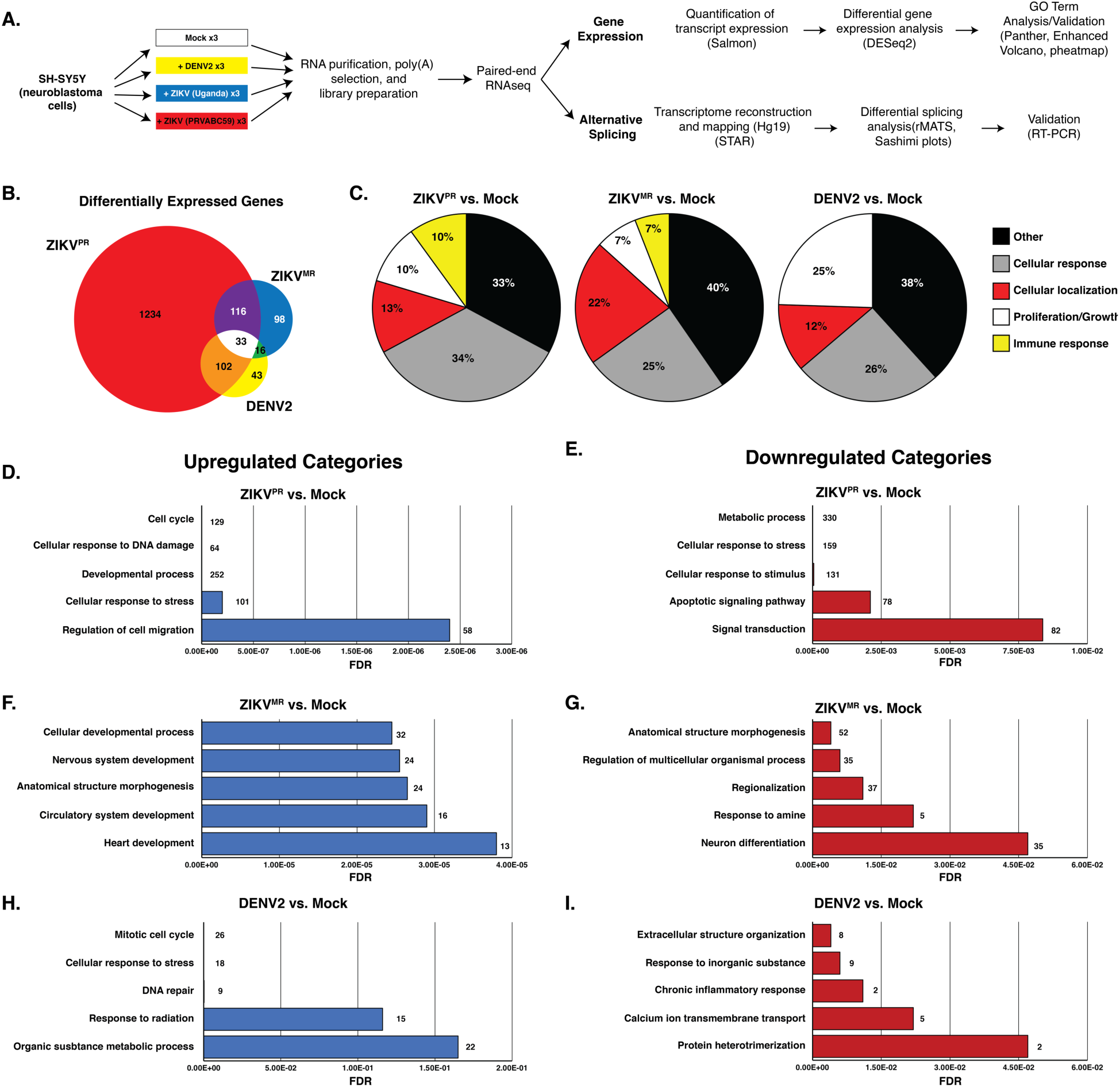
Transcriptome analysis of mock- and virus-infected SH-SY5Y cells. A) Schematic showing the pipeline from cells to differential gene expression and alternative splicing analysis. B) Venn diagram of differentially expressed (DE) transcripts between different viruses versus mock-infected SH-SY5Y cells. DE transcripts from ZIKV^PR^ versus mock, ZIKV^MR^ versus mock, DENV2 versus mock are highlighted in the red, yellow and blue circles respectively. C) All differentially expressed genes for each condition were input into ShinyGO(2.0) and the top 25 biological GO terms were categorized into five types and the distribution of each type presented in a pie chart. D, F, H) Top five functional categories derived from statistically significant upregulated genes for the indicated condition. E, G, I) Top five functional categories derived from statistically significant downregulated genes for the indicated condition. GO terms are annotated on the Y-axes, and false discovery rates (FDR) are represented on the X-axes.

To determine the gene categories broadly affected by these viruses, gene ontology (GO)-term analysis was performed on all statistically significant DE genes for each virus compared to mock. The top 25 GO-terms were categorized into five different types (cellular response, cellular localization, cell proliferation/growth, immune response and other), and plotted as pie graphs (Figure 2C). Interestingly, ZIKV^PR^ induced a greater percent of genes related to cellular response compared to ZIKV^MR^ and DENV (Figure 2C), while ZIKV^MR^ affected cellular genes influencing cellular localization (Figure 2C). Additionally, DENV2-altered genes were more heavily linked to cell proliferation and growth compared to either ZIKV strain. Notably, only ZIKV-infected cells had GO terms associated with the immune response (Figure 2C). In SH-SY5Y cells infected with ZIKV^MR^, 6% of DE genes were categorized under immune response, while 10% of genes DE in cells infected with ZIKV^PR^ were immune response genes (Figure 2C).

We also examined the top 10 DE genes from each infection. Notably, the top DE genes in ZIKV^PR^ infection were all upregulated, while in ZIKV^MR^ and DENV2 infected cells the top 10 DE genes were up- and down-regulated (Figure S2). Moreover, among the top 10 DE genes we observed little-to-no overlap between the viruses (Figure S2). Indeed, only *HSPA5* abundance was dramatically and modestly upregulated in ZIKV^PR^ and ZIKV^MR^-infected cells, respectively. Three (*ATF3, DDIT3* and *HSPA5*) of the top 10 genes upregulated in ZIKV^PR^ are involved in the stress response, and two genes were associated with lipid metabolism (*LDLR* and *SREBF1*). Interestingly, three of the upregulated genes in ZIKV^PR^ were non-coding RNAs (*SNHG15, SNHG17* and *OLMALINC*) (Figure S2). GO term analysis outside the top 10, revealed a large number of genes related to ER stress, protein misfolding, and PERK-mediated apoptosis (data not shown).

To examine possible links among the top 10 DE genes between each virus- and mock-infected cells, the top 10 DE genes were input into STRING, an online platform that is part of the ELIXIR infrastructure [52]. Connections between different proteins were only found when comparing ZIKV^PR^ to mock-infected cells (Supplemental Figure 2F) and these connected nodes are involved in PERK-mediated apoptosis (Supplemental Figure 2F). For example, apoptotic activation by PERK occurs when ATF4/ATF3 complex activates DDIT3, which in turn activates GADD34 [53]. These data suggest that ZIKV^PR^ impacts expression of genes associated with ER stress at multiple nodes.

We next performed GO analysis on genes that were differentially up- or downregulated during virus infection. Consistent with earlier transcriptomic studies, ZIKV^PR^ modulated the levels of genes associated with cell cycle, cellular response to DNA damage and stress, and apoptosis (Figure 2D and 2E). Indeed, the “cellular response to stress” was overrepresented in both up- and downregulated genes for ZIKV^PR^-infected cells (Figure 2D and 2G). Of the top 5 upregulated GO terms, only ZIKV-infected cells had terms linked to developmental processes (Figure 2D-2G), while those genes upregulated in DENV2-infected cells were linked to terms associated with cell response, including response to stress, radiation and inorganic substance (Figure 2H and 2I).

### ZIKV^PR^ notably upregulates immune response genes compared to ZIKV^MR^ and DENV2

Immunofluorescence and viral RNA abundance showed that SH-SY5Y cells were similarly infected by ZIKV^PR^, ZIKV^MR^ and DENV2 (Figure 1). To further understand the biological effect of each virus in SH-SY5Y cells, we analyzed the DE expression and used GO analysis to predict effects on cellular processes. We observed that 651 and 703 genes were up- and down-regulated respectively following ZIKV^PR^ infection (Figure 3A). In contrast, only 69 and 106, and 47 and 66 genes were up- and downregulated following ZIKV^MR^ and DENV2 infection, respectively (Figure 3A). This differential gene analysis showed only two genes (*HECW2* and *SATB1*) that were upregulated following infection with all three viruses, versus nine common genes (*ACHE, PIEZO1, ENO3, CDHR1, CCDC24, VCAM1, SLIT1, C4A*, and *ZNF321P*) that were downregulated (Figure 3A). While 20 upregulated genes were common between the ZIKV isolates, and 34 genes were similarly upregulated between ZIKV^PR^ and DENV2, ZIKV^MR^ and DENV2 shared no common genes (Figure 3A). A similar paired comparison between the viruses revealed common downregulated genes (Figure 3B).

**Figure 3:**
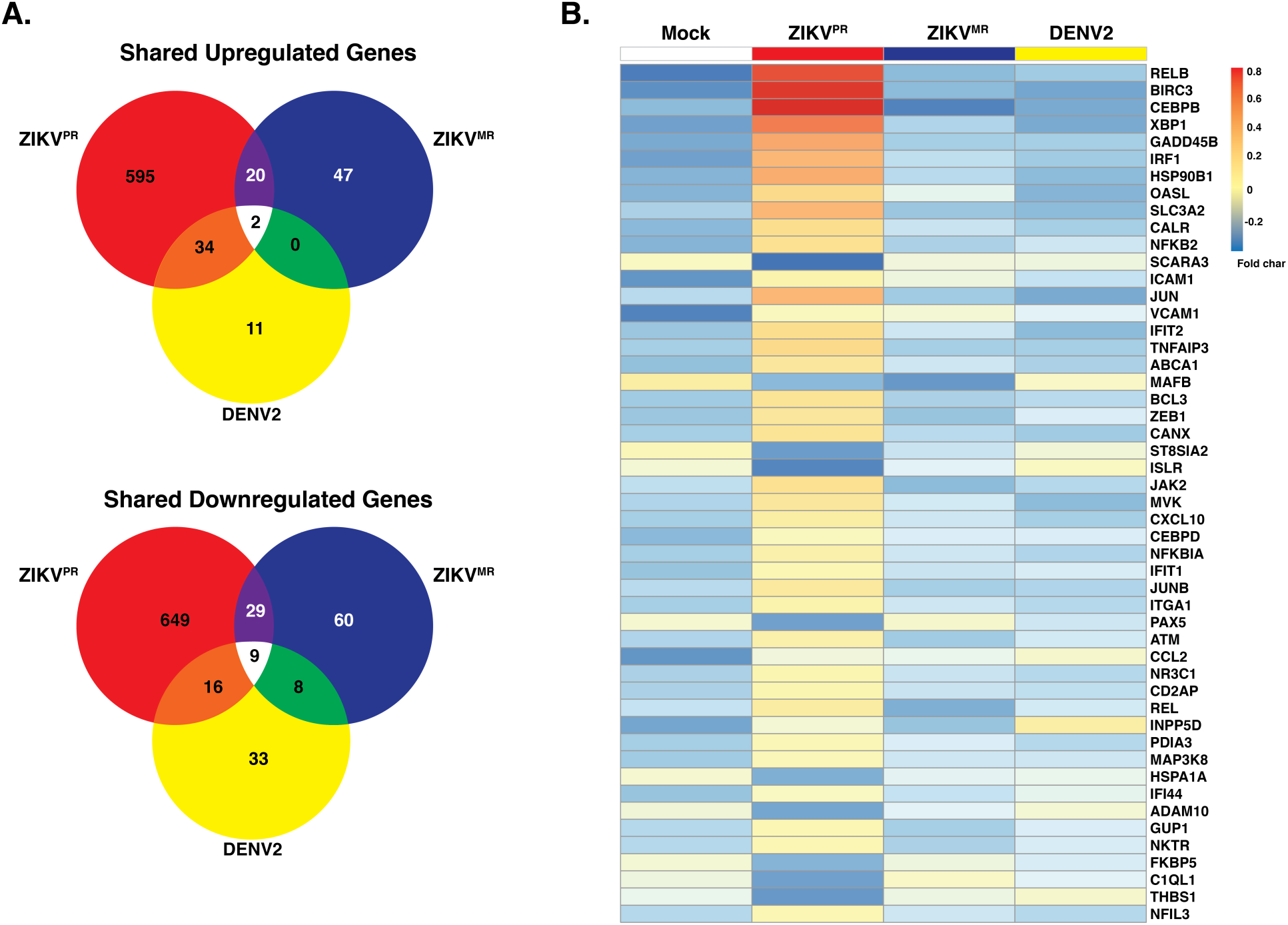
Upregulation and downregulation of genes in SH-SY5Y cells infected with ZIKV^PR^, ZIKV^MR^ and DENV2. A) Venn diagram of differentially expressed genes between infected SH-SY5Y cells that were differentially upregulated (top) or downregulated (bottom). DE transcripts from ZIKV^PR^, ZIKV^MR^, and DENV2 are within the respective red, yellow and blue circles. B) Heatmap of the top 50 statistically significant differentially expressed genes in ZIKV^PR^-infected cells compared to mock that are involved in the immune response. Three replicates from each condition were collapsed and normalized to mock. The color code shows the fold-change of the immune response genes in each virus infection.

Analysis of the top 25 GO terms in SH-SY5Y cells infected with the ZIKV isolates revealed an effect on immune response genes (Figure 2C). We therefore examine the expression of 52 genes that were categorized as immune response between mock and the three viruses (Figure 3B). Overall, we noted significant changes in gene expression following ZIKV^PR^ infection, compared to mock, ZIKV^MR^ and DENV2 (Figure 3B). Twenty percent of these immune response genes were downregulated following ZIKV^PR^ (Figure 3B). *RELB, BIRC3, CEBPB* and *XBP1* were greatly upregulated in ZIKV^PR^-infected cells (Figure 3B). Innate immune response genes such as *IRF1, OAS1, IFIT1* and *IFIT2*, were also all upregulated in ZIKV^PR^ infected cells (Figure 3B and Figure S3).

*MAFB, ST8SIA2, ISLR, HSPA1A*, and *ADAM19* were downregulated in both ZIKV-infected cells (Figure 3B). This subset of genes is interesting because of the association with developmental processes. For example, *ADAM19* is known to participate in neuromuscular junction formation [54], while *MAFB* loss-of-function and dominant-negative mutations result in a congenital eye-movement disorder known as Duane retraction syndrome [55]. Neuromuscular maldevelopment such as arthrogryposis was widely correlated to Zika congenital syndrome [10], and more than 80% of the infants with microcephaly born from ZIKV-infected mothers had ophthalmoscopic abnormalities [10]. Interestingly, *ADAM19* and *MAFB* were both upregulated in DENV-infected cells (Figure 3B).

In ZIKV-infected cells, *ICAM1* and *VCAM1* were upregulated, but were downregulated in DENV2-infected cells (Figure 3B). *ICAM1* and *VCAM1* are involved in immune cell migration across the blood-brain barrier [56,57], where an increase in immune cells in the brain might contribute to the neuropathogenesis associated with intrauterine ZIKV infection. ZIKV and other flaviviruses heavily depend on the secretory pathway for maturation of viral progeny. It is therefore interesting that of the 50 significantly altered genes associated with immune response, 21 contained endoplasmic reticulum (ER)-related, Golgi-related, and/or plasma-membrane-related GO cellular component terms within the top 5. These findings show that ZIKV^PR^-induced changes in gene expression significantly alter levels of genes involved in both the immune response and proper cellular development.

When we compared ZIKV^PR^ versus DENV2 DE genes, 138 genes associated with apoptotic signaling were downregulated (Figure S2E). Similarly, 23 apoptotic signaling pathway genes were downregulated when comparing ZIKV^PR^ to ZIKV^MR^, indicating that the modern isolate of ZIKV has developed strategies to usurp antiviral pathways of acquired mutations allowing for the suppression of the host immune response (Figure S2D). Interestingly, ZIKV^MR^ infection resulted in an upregulation of apoptotic genes when compared to DENV2 (Figure S3C).

### Exon skipping is a major splicing event during ZIKV^PR^, ZIKV^MR^ and DENV2 infection

While previous transcriptomic studies have been undertaken during ZIKV infection, we were interested in changes in alternative splicing (AS) induced following flaviviral infection. To this end we focused on five AS events (Figure 4A). Splicing of pre-mRNAs can be broken into five major categories, the most frequently observed event being skipped exon (SE) (Figure 4A). In addition to SE, alternative 5’ or 3 splice sites, mutually exclusive exons, and the retention of introns can also occur (Figure 4A). AS defects resulting in disease can occur when sequences on the pre-mRNA required for correct splicing are mutated or when regulatory factors essential for splicing are mutated [58]. These two scenarios can result in mis-splicing and thus a reduction in functional protein product or an imbalanced production of mature mRNA isoforms that contribute to disease.

**Figure 4:**
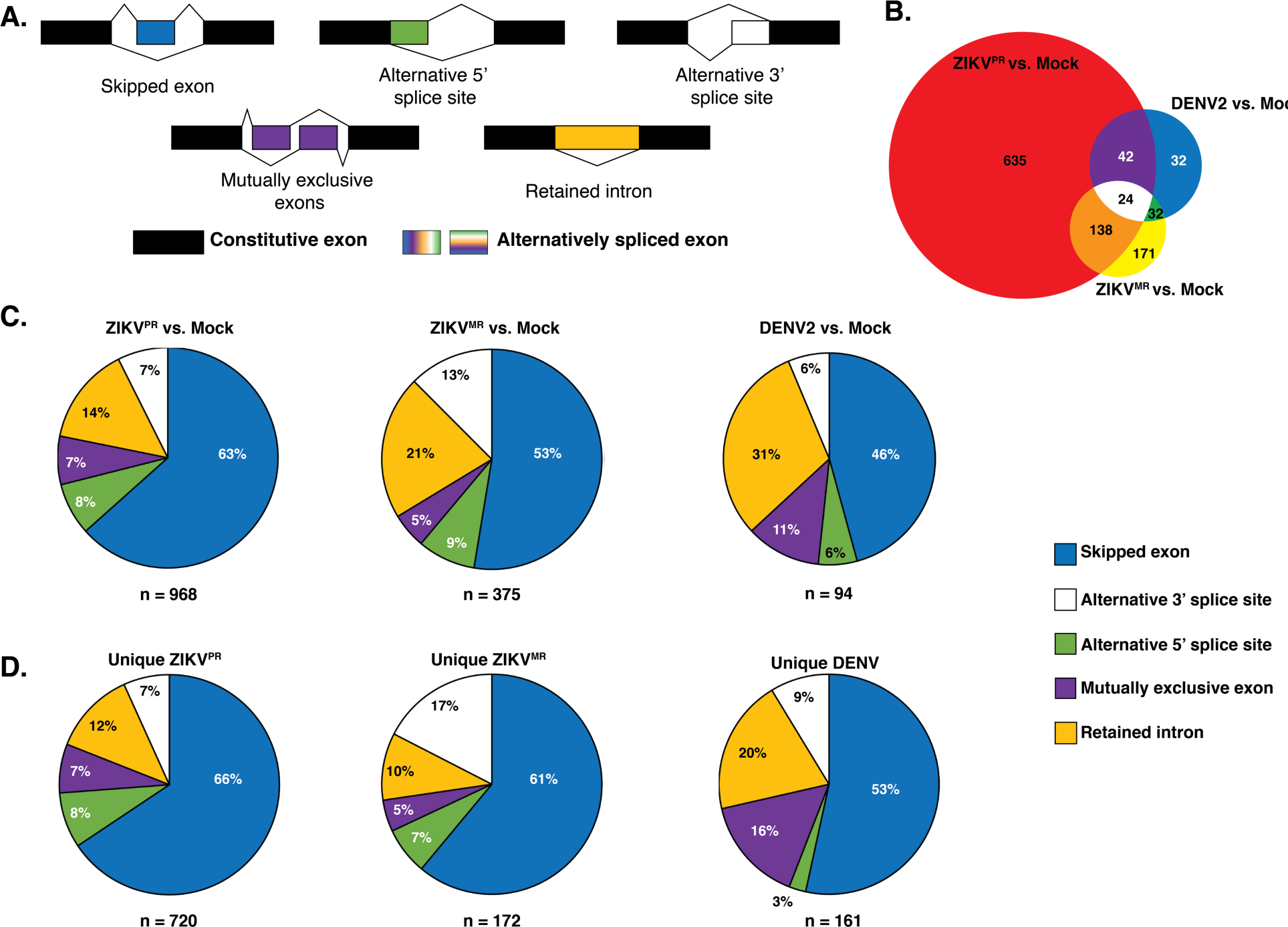
Analysis of alternative splicing events virus versus mock-infected SH-SY5Y cells. A) Schematic illustrating of the five alternative splice events characterized. The unchanged exons are black while the differentially spliced exons are color coded. B) Venn diagram of shared and unique alternative splicing events between virus and mock-infected cells. DE transcripts from ZIKV^PR^ versus mock, ZIKV^MR^ versus mock, DENV2 versus mock are highlighted in the red, yellow and blue circles respectively. C) Pie charts representing the percent of each type of AS event for the three viruses versus mock-infected SH-SY5Y cells. D) Pie charts representing the percent of each type of AS events unique to each virus infection in SH-SY5Y cells. The segment colors match the AS event illustrated in 4A.

To analyze splicing events, sequences were aligned to the reference human genome hg19 using STAR with ∼93% of reads aligning uniquely in each library. To analyze splice variants, rMATs was used, which generated a list of ∼77,000 (∼55,000 SE events) potential AS events occurring in all replicates. To reduce false-positive splicing events, all AS events with a false <u>d</u>iscovery rate (FDR) greater than 0.05 and a change in <u>p</u>ercent <u>s</u>pliced <u>i</u>n (ΔPSI) of less than 0.1 were omitted (∼500-1500 events depending on libraries).

We initially had expected a large portion of AS events between ZIKV^PR^- and ZIKV^MR^-infected cells would be similar. However, only 114 events were shared between both ZIKV isolates (Figure 4B). Similarly, 18 AS events were shared between ZIKV^PR^ and DENV2-infected cells (Figure 4B). Similar to the differential gene expression analysis, infection with the modern Asian-America ZIKV^PR^ isolate resulted in >2.5 times more AS events compared to cells infected with the ZIKV^MR^ and DENV2 (Figure 4C). When analyzing the types of AS events between each virus compared to mock, we observed skipped exons accounted for 63%, 53%, and 46% of all significant AS events for ZIKV^PR^, ZIKV^MR^, and DENV2, respectively (Figure 4C). The second most prevalent AS event for all infections compared to mock was intron retention (Figure 4C). In DENV2-infected cells, over 30% of AS events were retained introns, while this AS splicing event was lower, 21% and 14% respectively, in ZIKV^MR^ and ZIKV^PR^ infected cells (Figure 4B). Interestingly, AS events unique to DENV infection revealed more mutually exclusive exon splicing (16%) compared to 7% and 5% in ZIKV^PR^ and ZIKV^MR^ respectively (Figure 4C).

To further understand the biological processes modulated by AS and virus infection, all SE events for each condition were analyzed for functional category enrichment (Figure S5A-C). Of the top 10 statistically significant functional categories enriched in cells infected with ZIKV^PR^, four were involved in splicing supporting the idea that the global splicing landscape was being dramatically altered (Figure S5A). A second commonality between the top 10 functional categories related to disassembly of various cellular components including organelles, ribonucleoprotein complex, and ribosomes (Figure S5A-C). It is possible that the modern Asian-American ZIKV isolate promotes the disassembly of cellular factors required for completing the viral life cycle. Finally, we observed a significant enrichment in AS genes associated with central nervous system myelination (Figure S5A). When analyzing the AS events during infection with ZIKV^MR^, only eight GO functional categories were statistically significant (Figure S5B). Of these, three related to protein localization, two of which are involved in nuclear import and export (Figure S5B). Nine significantly enriched categories were found in DENV2-infected cells (Figure 5C). When comparing all significant categories between viral infections, similarities were only found between ZIKV^MR^ and DENV2-infected cells, which included monosaccharide transport, cellular response to nitrogen compound, and regulation of establishment of protein localization (Figure S5B and S5C). Moreover, the only significantly enriched functional category strictly specific to the central nervous system was found in cells infected with ZIKV^PR^ (Figure S5A).

**Figure 5:**
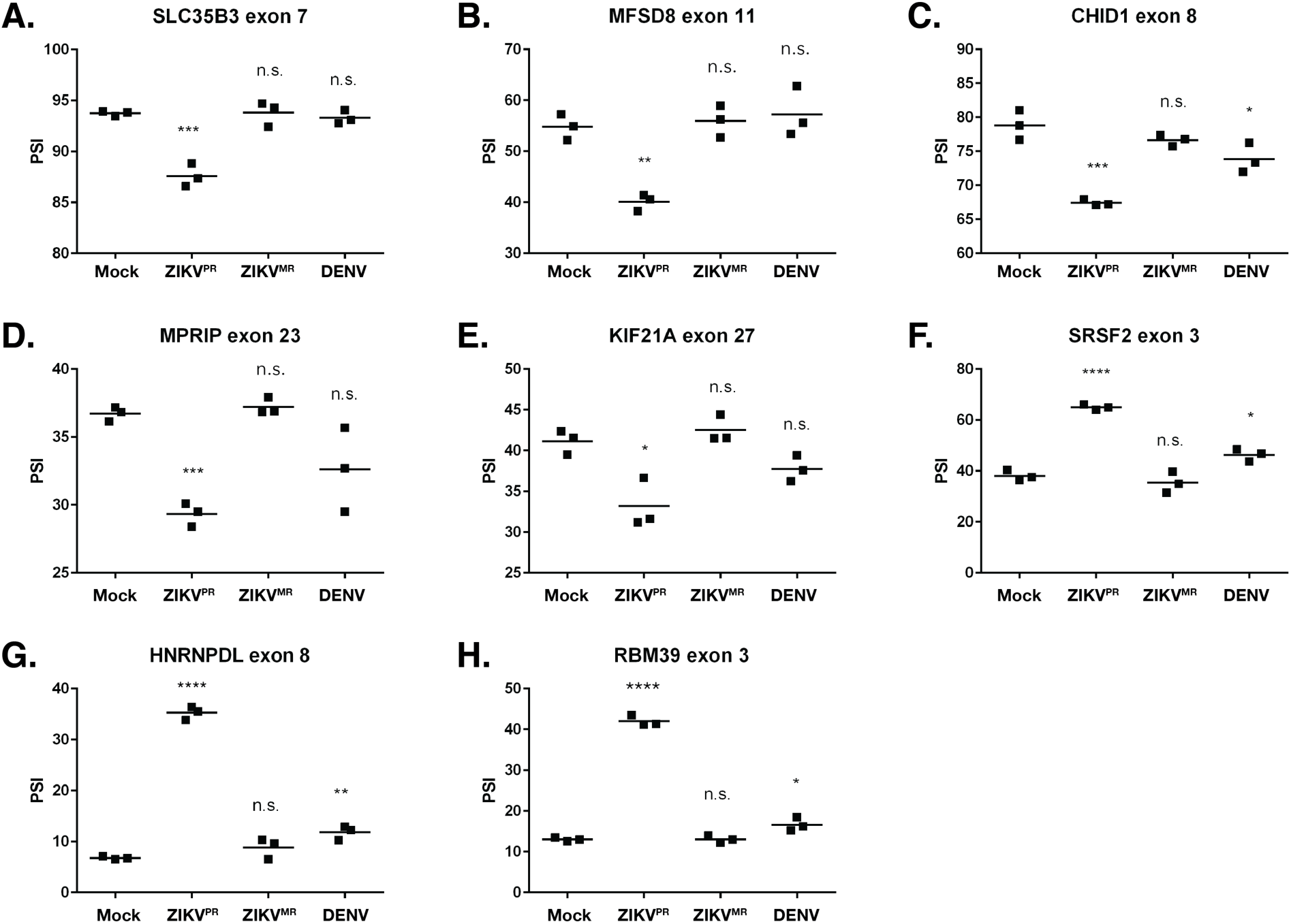
Validation of select alternative splicing events in mock-, ZIKV^PR^-, ZIKV^MR^- and DENV2-infected SH-SY5Y cells. A–I) RNA from each of three biological replicates were used for RT-PCR for the indicated gene. The percent spliced in (PSI) was calculated as described in the Materials and Methods. PSI indicates the change in inclusion or exclusion. Significance of the data were determined from the three independent experiments using the GraphPad software and performing an unpaired Student’s t-test; n.s. defines not significant, * P < 0.05, ** P < 0.01, *** P < 0.001, and **** P < 0.0001.

We were interested in which genes were specifically misspliced in each infection. Upon separating all AS events specific to each viral condition, we observed 968 genes were alternatively spliced in ZIKV^PR^-infected cells, compared to 375 and 94 for ZIKV^MR^- and DENV2-infected cells, respectively (Figure 4). To ascertain whether certain cellular pathways were preferentially being misspliced, all AS events for each type of splicing event were analyzed for enriched GO categories. Each of the top 10 enriched terms for ZIKV^PR^ specific AS events were linked to RNA processing with six of the 10 associated with splicing terms (Figure S5D). ZIKV^MR^ unique AS events were strongly linked to fatty acid metabolism and apoptotic pathways, while a majority of events resulting from DENV2 infection related to the nervous system (Figure S5E and S5F).

To validate AS events, we selected transcripts misspliced in ZIKV^PR^-infected cells compared to ZIKV^MR^ and DENV2. Exon skipping events comprised the majority of AS changes and can be easily assessed with RT-PCR. The genes chosen to validate the AS events were selected based on the lowest false discovery rate (FDR) and the highest read count from the RNA-seq data. RT-PCR was used to compare <u>p</u>ercent <u>s</u>pliced in (PSI) values for the chosen events. The same RNA that was used to perform RNA-seq was used in the alternative splicing RT-PCR assay. Primers were designed for each gene within the exons flanking the included/excluded exon (Figure S6). RT-PCR products were separated, and the intensity of the splicing products quantified (Figure S6). Thus, if virus infection promoted the inclusion of the exon, the PSI value increased. Conversely if the exon was excluded during virus infection, the PSI value decreased. Moreover, we have included Sashimi plots to visualize differences in the splice junctions of the selected transcripts from mock- and ZIKV^PR^ infected SH-SY5Y cell (Figure S6). The particular genes chosen for validation were: *SLC35B3, MFSD8, CHID1, MPRIP, KIF21A, SRSF2, HNRNPDL and RBM39*. These genes have interesting functionality. For example, *HNRNPDL* and *RBM39* function in transcription regulation [59,60], *HNRNPDL* and *SRSF2* regulate alternative splicing [59,61], *MPRIP* is involved in stress granule formation [62], *CHID1* has a role in pathogen sensing [63], and *KIF21A* has been implicated in neurological diseases [64]. Consistent with the selection criterion, RT-PCR validation of AS events showed significant missplicing in ZIKV^PR^-infected cells with the greatest difference in PSI values between Mock and ZIKV^PR^ was ∼28% (*RBM39*). For each of the genes examined we observed no significant splicing changes in the ZIKV^MR^-infected cells when compared to the mock infection. Interestingly, in DENV2-infectioned we determined small but significantly different changes of exon inclusion levels in *CHID1, SRSF2, HNRNPDL and RBM39*.

## Discussion

ZIKV infection during pregnancy has been linked many developmental and neurological abnormalities [12–15], and in adults Guillain-Barré syndrome is a consequence of ZIKV infection [16–19]. Although RNA-seq studies showed transcriptional dysregulation of genes associated with cell-cycle, DNA repair, immune response, cell death and microcephaly [21,22,25,26,65,66], these transcriptional changes were less dramatic between the African and Asian ZIKV strains [67]. These data suggested that transcriptional changes alone do not entirely explain ZIKV neuropathology, but rather other effects on mRNA such as cellular mRNA splicing and localization, and translation might contribute to ZIKV pathology. In this study we examined the consequence of ZIKV infection on alternative splicing. Specifically, we examined changes to the transcriptome and alternative splicing in a neuroblastoma cell line following infection with the modern Asian-American ZIKV isolate (ZIKV^PR^), the original African ZIKV isolate (ZIKV^MR^) and another mosquito-borne flavivirus not typically associated with neuropathies, namely dengue virus (DENV2). We show that infection with ZIKV^PR^ significantly changes the abundance of a high number of mRNAs compared to ZIKV^MR^ and DENV2, and that these cellular transcripts were associated with cellular processes such as cell cycle, cellular response to stress and DNA damage, and apoptosis. We additionally determined that ZIKV^PR^ dramatically alters alternative mRNA splicing, particularly that of exon skipping, compared to mock-, ZIKV^MR^- and DENV2-infected cells. Indeed, the observed changes in mRNA splicing support the premise that ZIKV affects cellular mRNA pathways beyond transcription and that understanding of changes to mRNA splicing might contribute to understanding the developmental and neurobiological anomalies associated with intrauterine ZIKV infection.

We observed a significant number of differentially expressed genes in SH-SY5Y cells associated with ZIKV^PR^ infection compared to ZIKV^MR^ or DENV2 infection. This result was unexpected particularly as our analysis of the percent infected cells and relative viral RNA abundance (Figure 1) between infection with the three viruses was similar. Therefore, our RNA-seq studies indicate that in SH-SY5Y cells, ZIKV^PR^ alters the transcriptional landscape more that either ZIKV^MR^ or DENV2 (Figure 2B). Zhang and colleagues [66], report 5- and 10-fold more differentially expressed genes following ZIKV^MR^ and DENV infection in human neural progenitor cells compared to our study in SH-SY5Y cells. The differences in the extent of transcriptional changes may be a result of virus infection in primary cells versus a cell line, and that we performed RNA seq studies following infection at a moi of 5, where at 24 hours post-infection only 40% of SH-SY5Y cells were infected (Figure 1 and Figure 2B).

Analysis of the initial transcriptomic data revealed within the top 10 differentially expressed genes in ZIKV^PR^-infected cells, a number of genes were involved in the unfolded protein response (UPR) or ER stress (Figure S2). These transcriptional changes are consistent with electron micrograph studies showing ultrastructural changes of the ER following ZIKV infection [68,69]. During the unfolded protein response, GRP78 (also known as HSPA5) releases IRE-1 α [70]. Consistent with studies investigating the unfolded protein response during ZIKV infection [71–73], we observed an upregulation of two genes within the unfolded protein response pathway (*HSPA5* and *XBP1*) in ZIKV^PR^-infected SH-SY5Y cells (Figure 3B and Figure S2).

ZIKV infection in different cells types induces the innate immune response. In primary human skin fibroblasts, the French Polynesia ZIKV isolate (ZIKV^FP^) was shown to promote the transcription of *RIG-I* (*DDX58*), *MDA5* (*IFIH1*), and *TLR3*, as well as interferon stimulated genes such as *OAS2, ISG15*, and *MX1* [74]. The 2’,5’-oligoadenylate synthase (OAS) and RNase L pathways have previously been shown to restrict flavivirus infection [75–77]. We were therefore surprised that *OAS3* was downregulated (Figure S4) while *OASL* was notably upregulated in ZIKV^PR^ infected cells (Figure 3B), suggesting that in neuronal cells, ZIKV gene expression might be restricted by another pathway such as by RIG-I (DDX58) and MDA5 (IFIH1). Indeed, In SH-SY5Y cells we also observed that *RIG-I* (*DDX58*) and *MDA5* (*IFIH1*) were upregulated by both ZIKV isolates, yet the mRNAs levels did not change during DENV2 infection (Figure S4). While TLR3 and TBK1 have been shown to impact ZIKV infection [13,22], where ZIKV^MR^ infection of human cerebral organoids reported an upregulation of TLR3 [13], and infection of neural stem cells by Mexican ZIKV isolates similarly altered the expression of immune response genes [78]. In ZIKV-infected SH-SY5Y cells however, we did not detect any changes in gene expression of *TLR3* and *TBK1* (Figure S4). The levels of the interferon receptor (IFNR) also did not change (Figure S4). Hanners and colleagues showed that in neural precursor cells that cytokines such as CXCL10 were not released following ZIKV^PR^ infection [69]. While we did not measure cytokine levels our RNA-seq data demonstrate that CXCL10 transcript levels are upregulated (Figure 3B). Additional differential expression analysis of immune-related genes revealed an upregulation of NF-kB inflammatory response genes including *BIRC3, NFKBIA, REL*, and *CXCL10* in ZIKV^PR^-infected cells (Figure 3C). These genes were similarly reported to be upregulated in ZIKV-infected human neural progenitor cells [79] and primary human skin fibroblasts infected with the French Polynesia isolate H/PF/2013 [74], although Simonin *et al*. [67] reported that in human primary neuronal cells infected with the Asian strain NF-kB genes were downregulated. Early transcriptomic studies demonstrated that during ZIKV infection the levels of known genes associated with microcephaly were altered. Disappointingly we did not observe changes in transcription of these microcephaly genes (data not shown). Our RNA-seq analyses in SH-SY5Y cells does not completely recapitulate the transcriptional landscapes in neural stem and progenitor cells, likely in part because SH-SY5Y cells are a neuroblastoma cell line derived from bone marrow [80,81]. It is noteworthy however that, SH-SY5Y cells can be differentiated into neurons [80], which presents an interesting opportunity to use these cells as a cell culture system to study ZIKV-associated Guillain-Barré syndrome in adults, a disease for which few studies have been undertaken.

To promote replication of the viral genome and evade the host innate immune response, viruses usurp numerous cellular pathways including altering host gene expression [82]. Indeed, viruses that replicate in the nucleus, such as herpes viruses, adenovirus and influenza A virus subvert the cellular RNA splicing machinery to broaden the coding capabilities of the viral genome [83–85]. Some viruses also alter splicing of cellular RNAs to promote the virus infectious cycle [82]. While flaviviruses do not splice their own RNA genomes, these positive-sense single-stranded RNA viruses have been shown to affect splicing of cellular transcripts [86]. In particular, the flavivirus RNA-dependent RNA polymerase (RdRp; NS5) is known to localize in the nucleus and bind to and interfere with mRNA splicing of multiple antiviral factors [86]. In our study ZIKV and DENV predominantly affect alternative splicing by skipping exons (Figure 4). Notably the number of AS events correlated with the number of genes differentially expressed following virus infection, where ZIKV^PR^ infection induced the highest number of AS events (Figure 4). Hu *et al*., [87] analyzed the RNA-seq data of ZIKV^MR^ infection of neural progenitor cells for alternative splicing events [21]. In this data set, the authors identified 262 AS events in 229 genes. Compared to mock-infected SH-SY5Y cells, we identified 375 and 968 AS events in ZIKV^MR^ and ZIKV^PR^-infected cells, respectively (Figure 4B). Interestingly, GO term analyses of the cellular processes impacted by ZIKV^PR^ infection revealed ten mRNA splicing-related terms (Figure S5A). Such categorization was absent in ZIKV^MR^ and DENV2-infected cells (Figure S5B and S5C). We validated eight AS events: *SLC35B6, MFSD8, CHID1, MPRIP, KIF21A, SRSF2, HNRNPDL*, and *RBM35*.

*SLC35B3* localizes primarily to the ER and Golgi and is a 3’-phophoadenosine 5’-phosphosulfate transporter involved in sulfation, a posttranslational modification associated with many molecules [88]. A close relative of SLC35B3, the CMP-sialic acid transporter (SLC35B2) has been linked to transcriptional regulation during acute inflammation [89]. The major facilitator superfamily (MFS) of proteins includes a number of transmembrane solute transporters that aid in the movement of small molecules across cell membranes [90]. MFS domain containing 8 (*MFSD8*), also known as *CLN7*, encodes for a putative lysosomal transporter ubiquitously expressed [91]. Mutations in CLN7 have been linked to a group of autosomal recessive neurodegenerative diseases known as neuronal ceroid lipofuscinoses (NCLs) [92]. Although several splice variants for MFSD8 are known to exist, the exact function or expression pattern of each are unknown. By altering the splicing of *SLC35B3* and *MFSD8* (Figure 5A and 5B), it is possible that ZIKV entry or exit, and thus the establishment of a successful, might be affected.

Very little is known about the native function of Chitinase Containing Domain protein 1 (*CHID1*). The gene for *CHID1* encodes a protein belonging to the superfamily glycoside hydrolase family 18 (GH18) suggesting it may play a role in carbohydrate binding [63]. Interestingly, RNA-seq gene expression profiles indicate CHID1 is expressed eight times higher in brain, almost 25 times higher in testis, and more than 20 times higher in ovaries compared to all genes in the respective tissue (Danielle Thierry-Mieg; Jean Thierry-Mieg, NCBI/NLM/NIH. <u>“AceView: Gene:CHID1, a comprehensive annotation of human, mouse and worm genes with mRNAs or ESTsAceView”</u>. Ncbi.nlm.nih.gov). *CHID1* has been shown to bind lipopolysaccharide (LPS) in a concentration-dependent manner suggesting this protein plays roles in pathogen sensing [63]. Perhaps the exon exclusion event carried out by ZIKV^PR^ (Figure 5C) specifically targets a domain required for pathogen sensing, or similar to *SLC35B3* and *MFSD8*, alternative splicing of *CHID1* could affect cellular components important for ZIKV entry or egress.

Myosin phosphatase-Rho interacting protein (MPRIP) targets myosin phosphatase to actin cytoskeleton and is required for the regulation of actin by RhoA and ROCK1 [93]. Previously, MPRIP was shown to regulate the formation of stress granules as overexpression of MPRIP led to the disassembly of stress granules in neuronal cells [94]. Interestingly, our transcriptomic studies indicated an increased expression of *MPRIP* in ZIKV^PR^-infected cells only (data not shown). Infection with ZIKV^PR^ leads to increased exclusion of exon 23 of MPRIP (Figure 5D). We and others have previously shown ZIKV disrupts the formation of arsenite induced stress granules (SGs) in non-neuronal cells [27,95]. Furthermore, we have shown that SG components exhibit both pro- and antiviral functions [27]. While the exact composition of neuronal SGs including those in SH-SY5Y cells is not known, perhaps the expression of alternatively spliced MPRIP differentially affects the formation of stress granules to impact ZIKV gene expression. KIF21A is a member of the KIF4 subfamily of kinesin-like motor proteins and is important for neural development in that it regulates microtubule dynamics and increased levels of it causes changes in axon morphology that cause axon guidance morphologies [64]. Hu *et al*., did not describe this splicing event, although it is interesting to note that in ZIKV^MR^ infection of cortical neural progenitor cells that MAP2, microtubule associated protein 2, was described to be alternatively spliced and could also affect axon morphology. ZIKV^PR^ leads to increased exclusion of exon 27 in *KIF21A* (Figure 5E). It would therefore be interesting to investigate the role of *KIF21A* with the inclusion of exon 27 during neurodevelopment and on ZIKV infection.

*SRSF2* belongs to the serine/arginine (SR)-rich family of pre-mRNA splicing factors. SRSF2 proteins constitute a large portion of the spliceosome and contain RNA recognition motifs (RRMs) RNA binding [96] and also play a role in the export of mRNA from the nucleus (Zhong XY, et al., 2009). Adenovirus, herpes simplex virus and vaccinia virus proteins have been shown to interact with SR proteins (Nilsson EC, et al., 2001; Kanopka A, et al., 1998) (Lindberg A and Kreivi JP, 2002; Hu B, et al., 2016), (Huang TS, et al., 2002). Of interest, *SRSF2, SRSF3, SRSF6*, and *SRSF7* were alternatively spliced in ZIKV^PR^-infected cells, but not in ZIKV^MR^-infected cells. Moreover, SRSF2 has been shown to suppress cassette exon inclusion in survival of motor neuron pre-mRNA (Moon H, et al., 2017). It is therefore possible that modulation of the *SRSF2* AS event (Figure 5F) contributes to the increased number of AS cases in ZIKV^PR^-infected cells of RNA.

*HNRNPDL* (JKTBP1) is a paralog of HNRNPD (AUF1), which has been shown to be important for transcription regulation and alternative splicing via the binding to 5’-ACUAGC-3’ RNA consensus sequences [59]. Interestingly, this consensus sequence is present a sequenced isolate of ZIKV from Brazil in 2015 (KU497555.1). The inclusion of exon 8 into hnRNPDL causes introduction of two exon junctions into the 3’ UTR of the second transcript. This inclusion results in a second junction located ∼60 nucleotides downstream of the natural termination codon suggesting that the hnRNPDL isoform including exon 8 is targeted for nonsense-mediated decay (NMD) [97]. In ZIKV^PR^ infected cells we confirmed a significant increase in the inclusion of exon 8 after infection with ZIKV^PR^ (Figure 5G), suggesting *HNRNPDL* could be a target for NMD, although this function would need to be verified. Interestingly, Hu *et al*., [87] also reported that HNRNPD was alternatively spliced in ZIKV^PR^ infected cortical neural progenitor cells.

RNA binding motif protein 39 (*RBM39*), a member of the U2AF65 protein family, is a splicing factor and transcriptional co-activator of AP-1/Jun and estrogen receptors [98,99]. Moreover, shRNA downregulation of RBM39 activity leads to decreased expression of cell-cycle progression regulators, inhibits protein synthesis, prevents the phosphorylation of c-Jun [100], and when complexed with TBX3 is necessary for preventing primary cell and mouse embryo senescence [101]. Interestingly, RBM39 mRNA expression is abundant in many immune system-associated cells raising the possibility that modern ZIKV isolates may target directing or indirectly immune related genes for alternative splicing by changing the splicing of *RBM39* (Figure 5H) [102].

The 2015 outbreak in the Americas, and in particular the devastating developmental and neurological defects in newborns catapulted ZIKV into the scientific limelight. To understand these virus-associated neuropathologies at a genetic level a number of transcriptomic studies were undertaken [12,20–23,25,26]. In this study we demonstrate that changes in gene expression are not the only effect ZIKV has on cellular RNA homeostasis, but that alternative splicing likely also has a role. Our analyses establish the groundwork to investigate the role of alternative splicing in other tissues critical in ZIKV pathogenesis. Moreover, future studies will elucidate the effect specific alternative splicing changes on neurodevelopment, the immune response and ZIKV infection. The transcriptome landscape and alternative splicing are two pathways contributing to cellular RNA homeostasis (or ribostasis) and further studies on mRNA stability, translational profiles and changes in the epitranscriptomic landscape will provide new and critical insights on ZIKV-host interactions in neurodevelopment. Manipulation of these virus-host interactions may eventually provide a foundation for developing new antivirals that will reign in this re-emerging and devastating virus.

## Author Contributions

Conceptualization, G.B. and C.T.P.; Methodology, G.B., C.S. and C.T.P.; RNA-seq analysis, R.M., G.B. and J.A.B.; Validation, G.B. and C.S.; Investigation, G.B., C.S. and R.M.; Writing – Original Draft Preparation, G.B. and C.T.P.; Writing – Review & Editing, G.B., R.M., C.S., J.A.B. and C.T.P.; Supervision, J.A.B. and C.T.P.; Funding Acquisition, C.T.P.

## Acknowledgements

We thank Dr. Brett Lindenbach (Yale School of Medicine), CDC, Dr. Laura Kramer (Wadsworth Center, NYSDOH) and Dr. Alexander Ciota (Wadsworth Center, NYSDOH) for the generous gifts of ZIKV stocks (Ugandan-isolate (MR766), and Puerto Rican-isolate (PRVABC59)) and DENV2. We gratefully acknowledge from Dr. Morgan Sammons (Department of Biological Sciences, University at Albany-SUNY) for assistance and advice with the RNA-seq experiments; and Dr. John Cleary for his thoughtful and insightful comments on the manuscript. We also thank members of the Pager and Berglund labs for valuable comments and suggestions on the manuscript. We are grateful to The Center for Functional Genomics, University at Albany-SUNY for the NextGen Sequencing services.

