## Supplemental Data for "Asian Zika virus isolate significantly changes the transcriptional profile and alternative RNA splicing events in a neuroblastoma cell line"


| | |
|---|---|
| Table S1. | List of primers used for RT-qPCR and validation of alternatively spliced mRNAs. |
| Figure S1. | Volcano plots of differential gene expression analysis. |
| Figure S2. | Analysis of the top 10 differentially expressed genes. |
| Figure S3. | Analysis of GO terms of upregulated and downregulated genes in virus-infected SH-SY5Y cells. |
| Figure S4. | Heatmap of differentially expressed interferon-related genes in mock- and virus infected SH-SY5Y cells. |
| Figure S5. | GO term analysis of alternatively spliced events in mock- and virus infected SH-SY5Y cells. |
| Figure S6. | Schematic, gel analysis of RT-PCR products and Sashimi plots of eight validated alternatively spliced targets. |

**Table S1: List of primers used for RT-qPCR and validation of alternatively spliced mRNAs**

| Primer target | Sequence (5' → 3') |
|---|---|
| $ZIKV^{MR}$ (FWD) | TTGGTCATGATACTGCTGATTGC |
| $ZIKV^{MR}$ (REV) | CAGGTCCCACCTGACATGC |
| $ZIKV^{PR}$ (FWD) | CCTTGGATTCTTGAACGAGGA |
| $ZIKV^{PR}$ (REV) | AGAGCTTCATTCTCCAGATCAA |
| DENV2 (FWD) | CAGGTTATGGCACTGTCACGAT |
| DENV2 (REV) | CCATCTGCAGCAACACCATCTC |
| β-Actin (FWD) | GTCACCGGAGTCCATCACG |
| β-Actin (REV) | GACCCAGATCATGTTTGAGACC |
| HNRNPDL (FWD) | CAACAGAGCACTTATGGCAAGG |
| HNRNPDL (REV) | CGTCCTGCAAGATGGGTTACT |
| SRSF2 (FWD) | AGGAGCGGTGTCCTCTTAAGA |
| SRSF2 (REV) | TTTTTCCCCAAGTCCTCCGTT |
| RBM39 (FWD) | CCGAACACGAGCACCACAG |
| RBM39 (REV) | GTTCTTCATGGCCGTTGGCA |
| MPRIP (FWD) | GATCCTGTGTCACCCGGCAA |
| MPRIP (REV) | CCGTTCTTGCACCGTCAG |
| KIF21A (FWD) | CCAGGCAGTCATCTCTATCAGA |
| KIF21A (REV) | GCACAGCTTTTGTATGCCCT |
| CHID1 (FWD) | GCTTCGTGGTGGAGGTCTG |
| CHID1 (REV) | GGGCCAGCTGCTCAAACTC |
| MFSD8 (FWD) | TACTTGCTGCTCTTGGGGT |
| MFSD8 (REV) | TTCCCCAAATGTGGTATTAGGG |
| SLC35B3 (FWD) | TGACAGCACAACTGCACCA |
| SLC35B3 (REV) | GCCTAATCCACTAGTGCATGT |

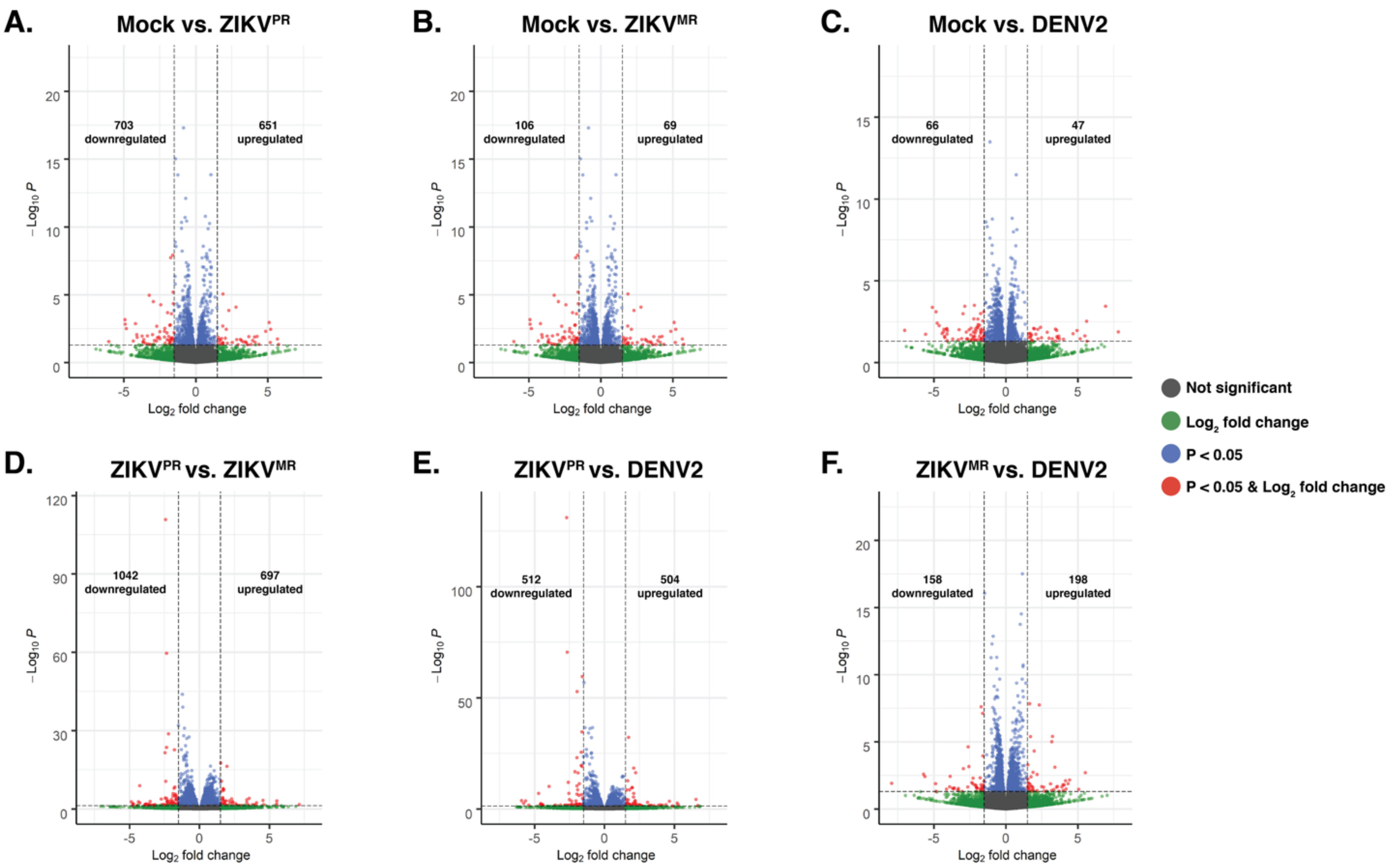


**Figure S1: Volcano plots of differential gene expression analysis in mock- and virus-infected SH-SY5Y cells.**
A–F) Volcano plots of differential gene expression analysis plotting the -$Log_{10}P$ versus $Log_2$ fold change for the indicated comparison. Values overlaid on each graph represent number of downregulated (left) and upregulated (right) genes for the indicated condition.

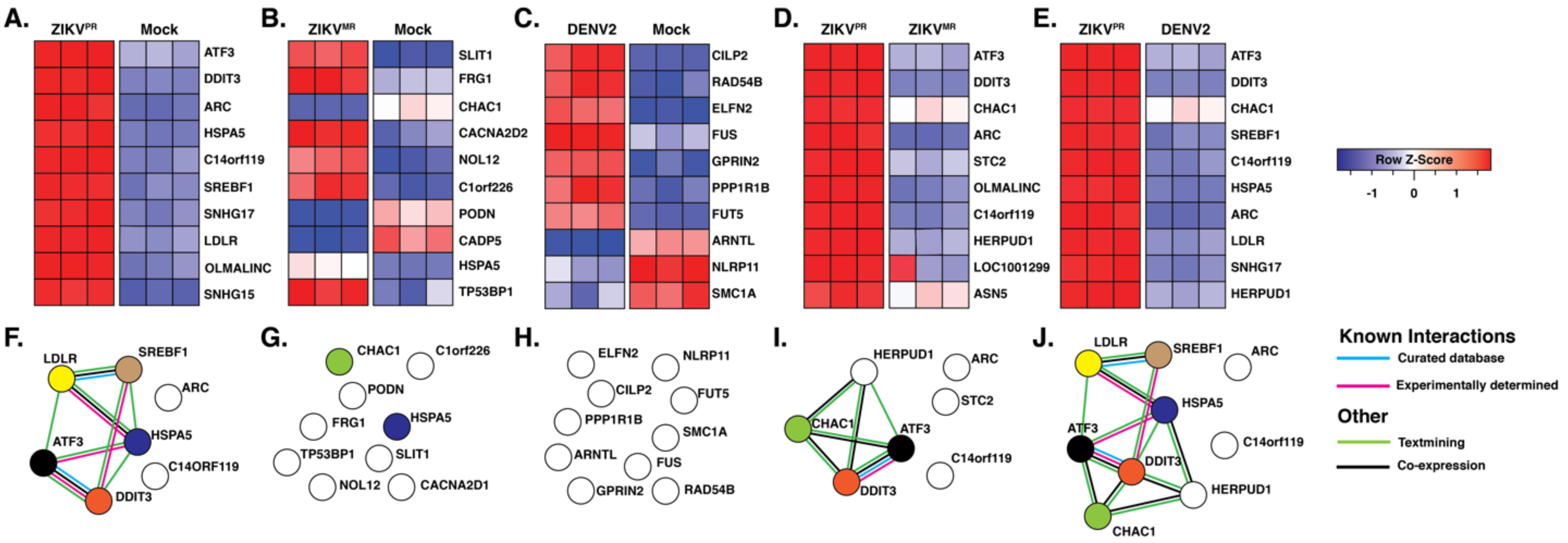


| ZIKV$^{PR}$ vs. Mock | ZIKV$^{MR}$ vs. Mock | DENV2 vs. Mock | ZIKV$^{PR}$ vs. ZIKV$^{MR}$ | ZIKV$^{PR}$ vs. DENV2 | |
|---|---|---|---|---|---|
| 7 | 9 | 10 | 7 | 9 | Number of nodes |
| 8 | 0 | 0 | 6 | 14 | Number of edges |
| 2.29 | 0 | 0 | 1.71 | 3.11 | Average node degree |
| 2.07E-07 | 1 | 1 | 1.37E-09 | 2.49E-14 | PPI enrichment p-value |

**Figure S2: Analysis of the top 10 differentially expressed genes.**
A-E) Heatmaps generated using Limma-voom. Shaded squares represent the number of standard deviations away from the mean (z-score) for each of three biological replicates. F-J) Known (blue and pink lines) and predicted (green and black lines) interactions between the top 10 differentially expressed genes of each comparison. Listed below node maps are the statistics generated from STRING.

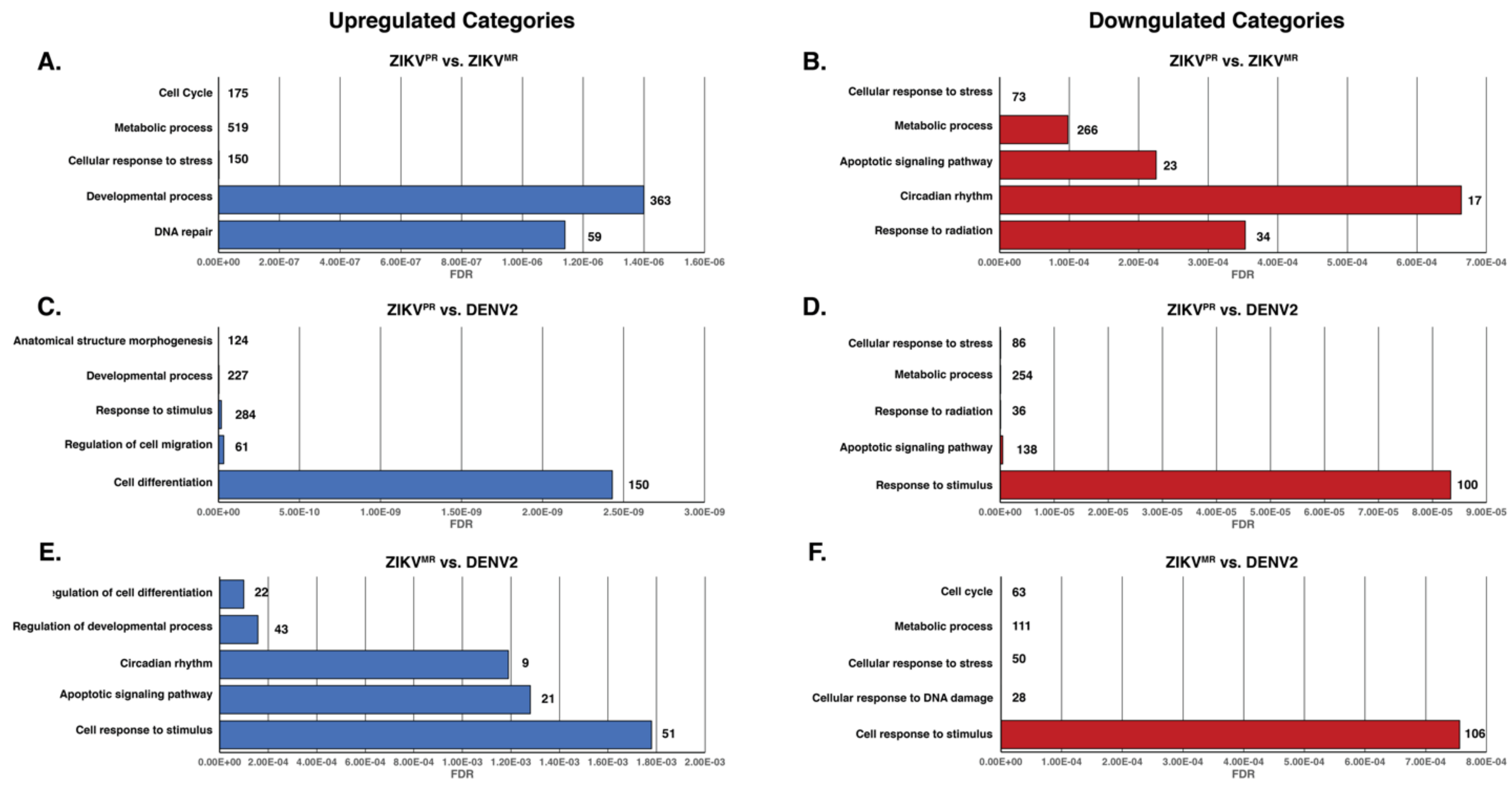


**Figure S3: Analysis of GO terms of upregulated and downregulated genes in virus-infected SH-SY5Y cells.**
All statistically significant genes that were differentially upregulated (blue) or downregulated (red) were input into enrichR. The top 5 categories were plotted. Numbers to right of bars indicate number of genes in particular category. GO terms are shown on the Y-axes, and false discovery rates (FDR) are represented on the X-axes.

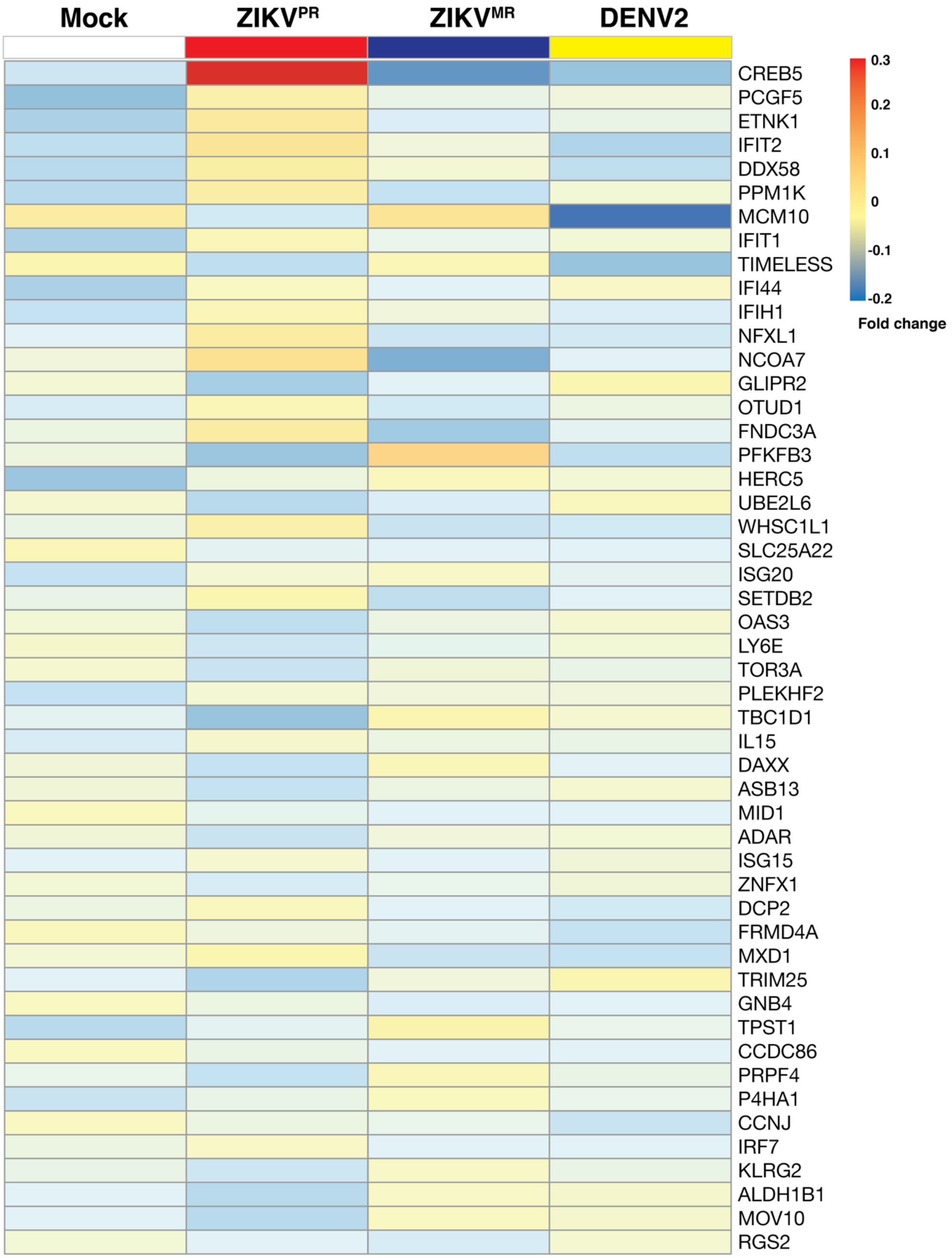


**Figure S4: Heatmap of differentially expressed interferon-related genes in mock- and virus infected SH-SY5Y cells.**

Heatmap of the top 50 differentially expressed genes in ZIKV$^{PR}$-infected cells compared to mock that are related to the interferon pathways. Bolded gene names are significant ($p < 0.05$). Three replicates from each condition were collapsed and normalized to mock.

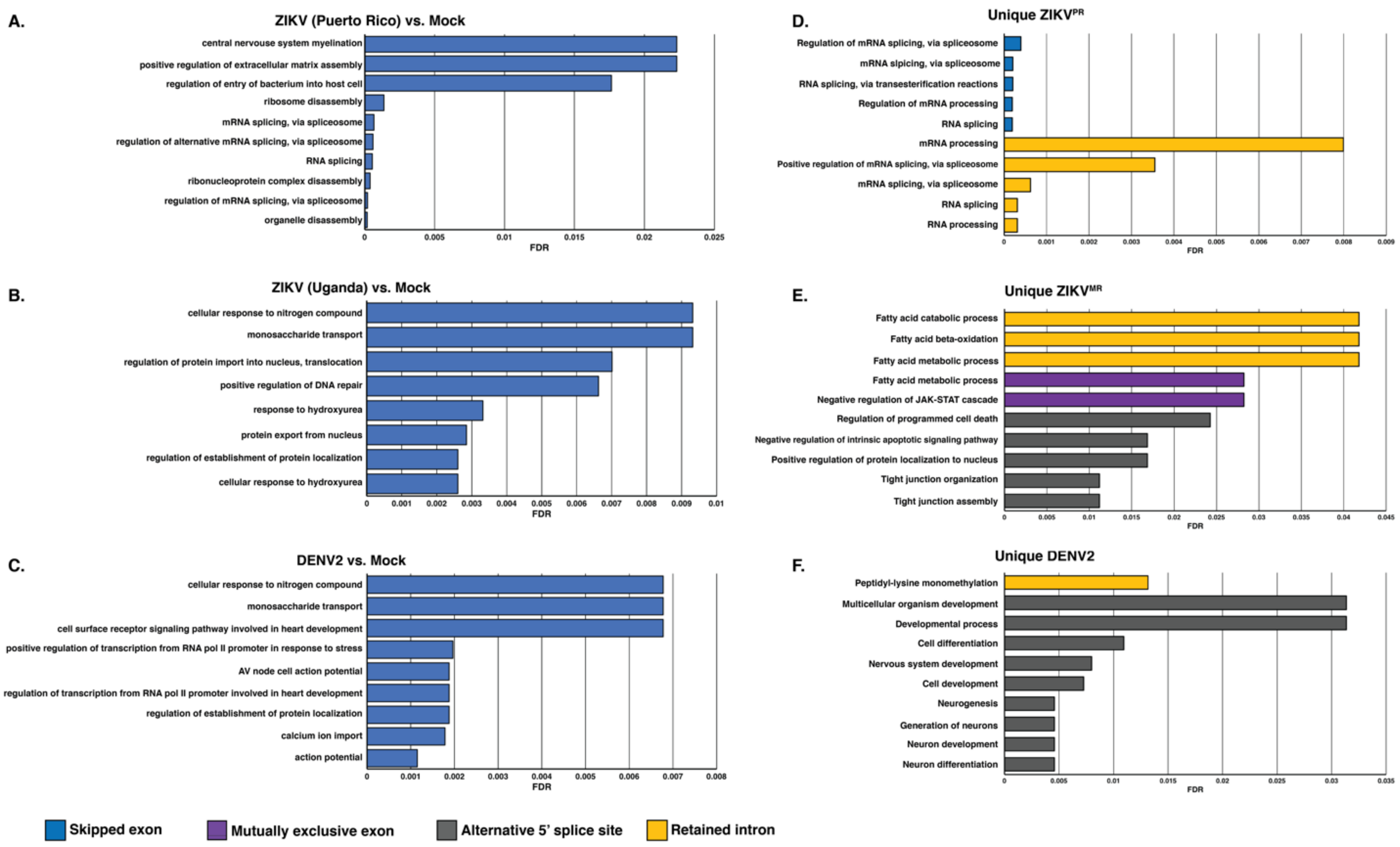


**Figure S5. GO term analysis of alternatively spliced events in mock- and virus infected SH-SY5Y cells.**
All gene IDs of a given subset were input into ShinyGO2.0 for functional enrichment. The top 10 functionally enriched categories were plotted. A, B, & C) Analyses of enrichment of cellular processes when comparing virus versus mock-infected SH-SY5Y cells. D, E, & F) GO categorization of virus-specific alternatively spliced events. GO terms are annotated on the Y-axes, and false discovery rates (FDR) are represented on the X-axes.

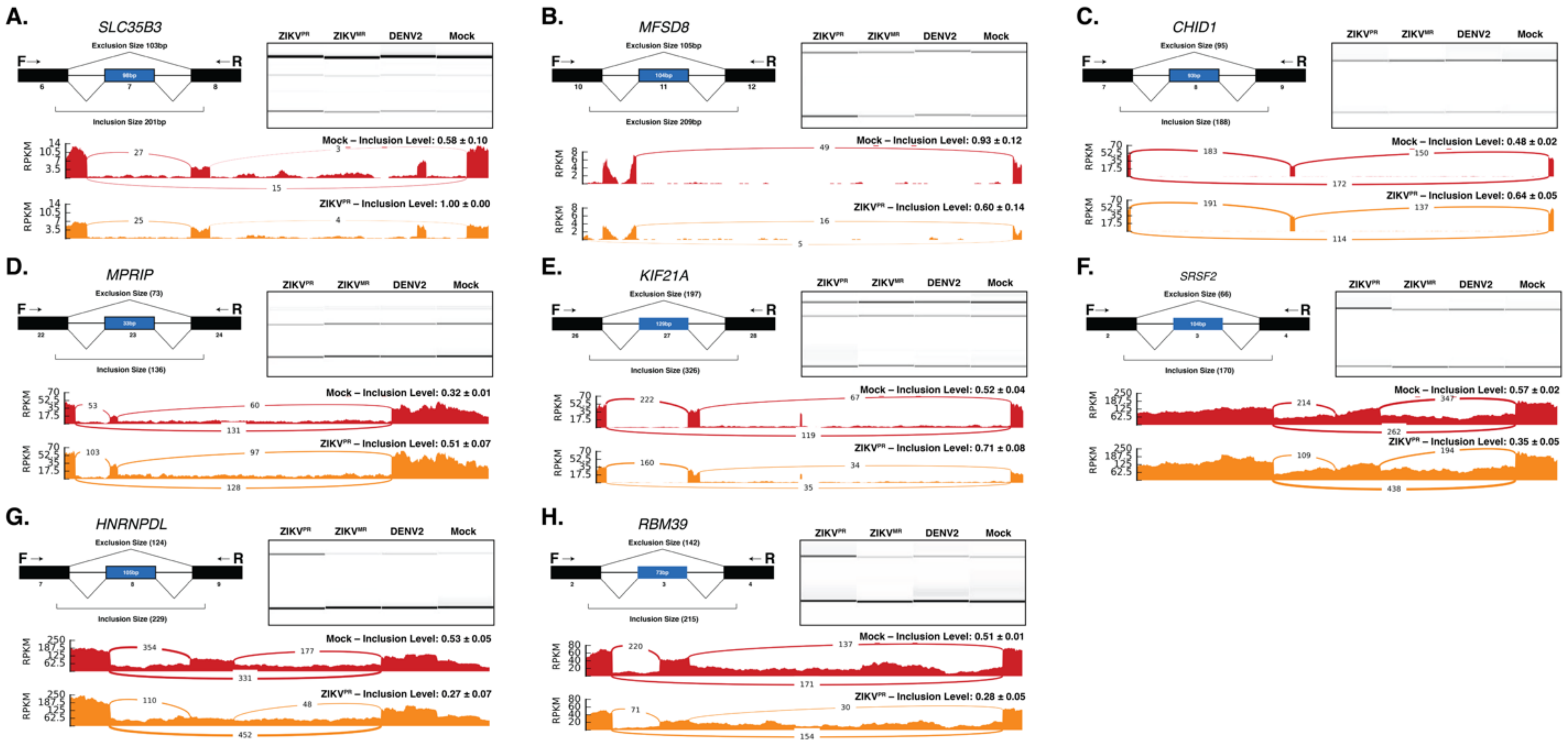


**Figure S6: Schematic, gel analysis of RT-PCR products and Sashimi plots of eight validated alternatively spliced targets.**
For each panel: (top left) a schematic illustration of constitutive exons (black) and the cassette exon (blue) of interest. Exclusion and inclusion size were predicted based off primer binding sites; (top right) representative gel image of alternative splicing analysis where top band represents inclusion of the cassette exon and the lower band represents exclusion of the cassette exon; (bottom) one representative Sashimi plot from Mock-infected and $\text{ZIKV}^{PR}$ infected SH-SY5Y cells, from each transcript validated. Values listed for inclusion levels represent the average of all three biological replicates ± standard deviation.
